# Meta-analysis of genetic mapping studies in mice reveals candidate epilepsy modifier genes that are outside the current drug development landscape

**DOI:** 10.1101/2025.08.15.669922

**Authors:** Giovanna L. Durante, Anna L. Tyler, Rod C. Scott, Amanda E. Hernan, J. Matthew Mahoney

**Affiliations:** The Jackson Laboratory, 600 Main St., Bar Harbor, ME 04609; Nemours Children’s Hospital, 1600 Rockland Rd., Wilmington, DE, 19803

**Author notes:** **FUNDING INFORMATION** This research was funded by the National Science Foundation Grant #2244034 (GD). **ETHICS STATEMENT** Data and software to recapitulate these analyses are publicly available and open source.

**Keywords:** Genome-wide association study, Systems biology, Multi-species analysis, Drug target prioritization

## Abstract

**Objective:** Despite decades of development in anti-seizure medications, approximately 30% of individuals remain refractory to all treatments, and none of the existing therapies are disease-modifying. Identifying targets outside the current preclinical paradigm is critically important. This study aimed to characterize the landscape of current epilepsy treatments at the level of gene interaction networks and identify novel genetic modifiers of epilepsy as potential novel therapeutic targets.

**Methods:** We performed a functional network analysis to score genes based on their interactions with known epilepsy genes and integrated these functional scores with population-genetic data and drug tractability information. In parallel, we performed a meta-analysis of genome wide association studies of epilepsy-related phenotypes in genetically diverse mice using a large compendium of historical phenotyping data. Genes within mapped loci were prioritized based on functional rankings and genomic evolutionary rate profiling (GERP) was used to identify highly SNPs at evolutionarily constrained positions.

**Results:** Functional network analyses of known epilepsy genes revealed a strong involvement of neurodevelopmental processes in epilepsy pathogenesis, which are not targeted by existing or emerging treatments. Meta-analysis of seizure traits in mice identified 118 non-overlapping loci harboring potential seizure phenotype modifiers. Using functional rankings, we prioritized 168 candidate genes within these loci and used GERP scores to filter down to 75 single-nucleotide polymorphisms as candidate variants within these genes. Among them, five genes—*Ephb2, En2, Cadps2, Igsf21*, and *Cep170*—contain regulatory variants in evolutionarily constrained sites. Four of these genes are validated as modifiers of neurological traits, including epilepsy susceptibility.

**Significance:** This study prioritized epilepsy modifier genes that are strongly predicted to influence neurodevelopmental processes that are underrepresented among current therapeutic targets. Furthermore, the identified genes represent novel candidate modifiers with potential clinical relevance. Our systems-level analysis offers a novel view into the potential target landscape, pointing toward promising new directions for disease-modifying treatments.

**KEY POINTS:**

- Systems-level analysis of human epilepsy genes reveals neurodevelopment, synaptic plasticity, and membrane excitability as key biological processes defining risk.
- The current drug-development landscape does not target neurodevelopmental processes
- Meta-analysis of mouse genome-wide association studies of epilepsy-related phenotypes reveal hundreds of loci with putative modifier genes.
- Prioritization of candidate genes reveals high-quality candidates that are known to influence neurodevelopment and synaptic plasticity, and be druggable.

## 1 INTRODUCTION

Epilepsy is a genetically complex family of debilitating neurological disorders characterized by recurrent, unprovoked seizures that affects over 50 million people worldwide.(1) Seizures are often attributed to an excitatory-inhibitory (EI) imbalance at the level of the neuronal membrane, and the mechanisms of action of many anti-seizure medications (ASMs) are construed in this framework, *e*.*g*., restoring inhibition by directly activating GABA receptors (2). However, given the complexity of neural circuits, the EI-imbalance framework belies a more nuanced picture of epilepsy as fundamentally a phenomenon of dynamic networks operating at multiple spatial and temporal scales (3). Critically, approximately 30% of patients fail to become seizure free with medication (4), underscoring the need for new therapeutic strategies. The heterogeneity of epilepsy, spanning diverse etiologies, seizure types and patient responses, further complicates treatment and necessitates a system-level understanding of the biological processes predisposing a brain to develop seizures in order to uncover robust, broadly effective therapeutic targets.(5)

Many forms of epilepsy are heritable and to date hundreds of epilepsy-associated genes have been identified.(6) While rare monogenic syndromes provide valuable mechanistic insight,(7) the majority of patients have idiopathic disease caused by some combination of polygenic risk and environmental exposure.(8) Importantly, genetic variation offers a powerful lens for understanding disease mechanisms, as germline variants can influence seizure susceptibility without being confounded by disease progression. In the context of drug development it has been observed that genetic-trait associations for a drug target dramatically increases the likelihood of clinical trial success.(9) However, despite many genetic associations, breakthrough therapies for epilepsy have not emerged to harness these mechanisms and significantly reduce overall disease burden. Part of the difficulty is rigorously prioritizing among many potential therapeutic targets to find the robust mechanisms of resilience that can induce a brain to a healthy state.

Animal models, particularly genetically diverse mouse populations, offer a complementary strategy to dissect the genetics of epilepsy. In mice it is possible to tightly control the etiology and timing of seizure insults to probe for genetic modifiers that mitigate against poor outcomes. Such modifiers represent highly attractive therapeutic targets, provided their mechanisms are conserved between mice and humans. Evidence from system-level analyses of human and mouse genetic and gene expression data strongly implicate shared genetic pathways driving epilepsy,(10) suggesting a high degree of translatability between species. Like in humans, genetic variability in epilepsy-associated phenotypes in mice has been widely observed. Heritable differences in induced and spontaneous seizure traits have been extensively studied in genetically diverse mouse panels, including the BXD and AXB/BXA recombinant inbred panels and in inbred strain surveys.(11) Legacy data from several of these studies have been cataloged in the Gene Network database,(12) making them available for re-analysis, providing an opportunity to identify broadly acting genetic modifier loci that expose key drivers of seizure-associated gene networks.

Here, we present a two-stage, cross-species systems genetics approach to prioritize novel candidate genes as potential disease-modifying drug targets for epilepsy. First, we construct an epilepsy interactome by integrating known epilepsy genes with human and mouse gene networks to identify conserved pathways and their overlap with existing drug targets. Second, we performed a meta-analysis of genetic mapping studies of seizure phenotypes in mice to prioritize seizure-modifier genes within the epilepsy interactome as potential disease-modifying drug targets. Our innovative framework is outlined in Fig. 1. This integrative, multi-species approach leverages the independent strengths of human association data and intervention studies across mouse genetic diversity to characterize novel genes and pathways involved in epilepsy. This work aims to advance the discovery of broadly effective, genetically informed therapies for epilepsy, addressing a critical unmet need in the field.

**Fig. 1.**
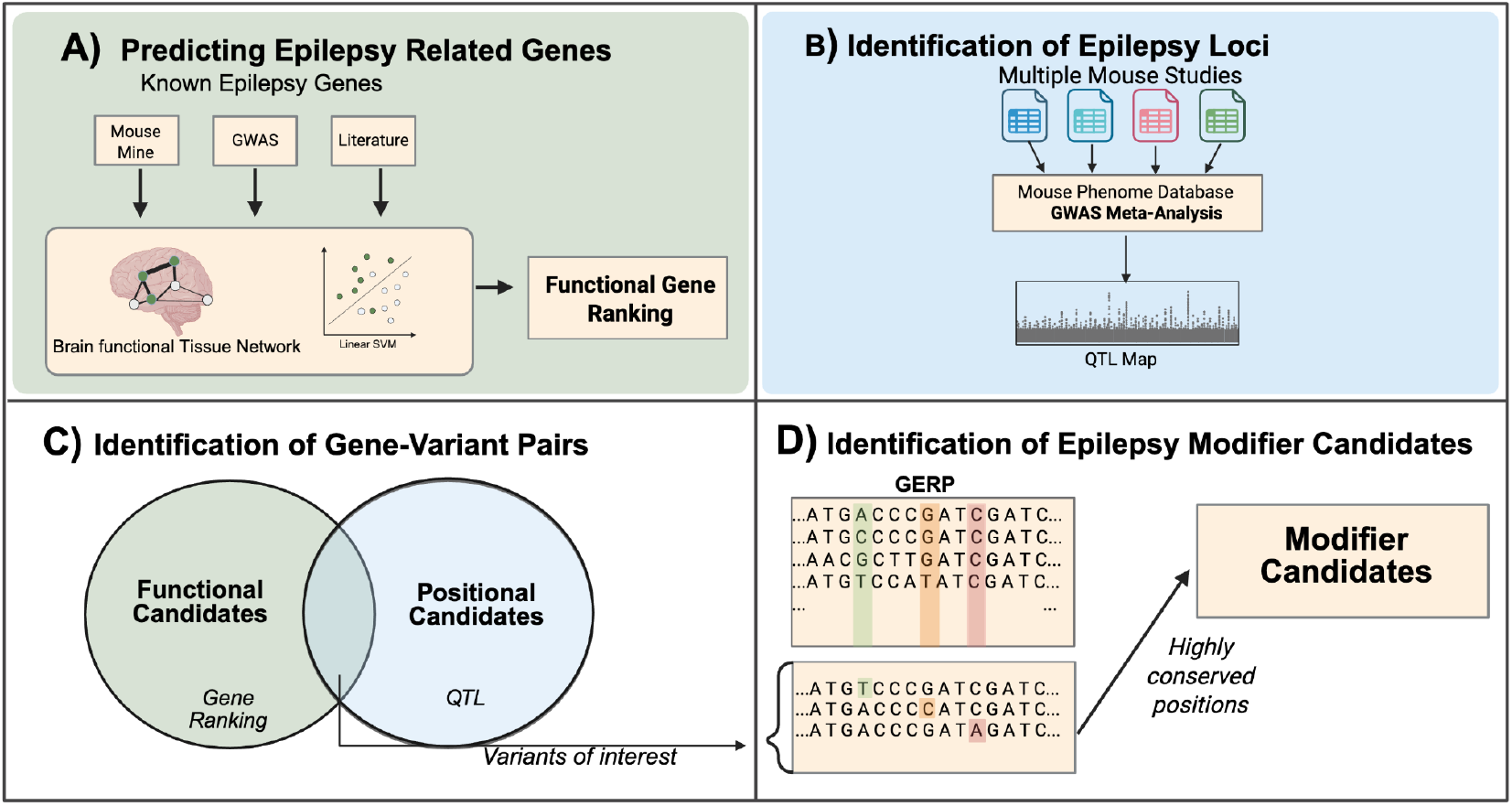
Workflow for integrating human and mouse genetic mapping data to prioritize modifier genes and variants. **(A)** We performed a network-based analysis of known epilepsy genes from multiple sources. We trained an ensemble of support vector machines (SVMs) to functionally score genes for involvement in epilepsy pathways based on connectivity within mouse and human brain functional networks. **(B)** Next, we performed a GWAS meta-analysis in mice to identify epilepsy-related loci containing modifiers of seizure phenotypes. **(C)** We prioritized candidate modifiers using positional and function information from (A) and (B) to produce a set of high-quality gene candidates. **(D)** To prioritize potential causal variants within high-quality candidate genes, we filtered variants based on GERP conservation score associated with the location of the variant. Figure made with Biorender.

## 2 MATERIALS AND METHODS

### 2.1 Network-based machine learning for functional gene prioritization

To gain a systems-level understanding of the interactions of genes involved in epilepsy, we applied a previously published network-based machine learning approach.(13) Briefly, we trained an ensemble of 100 support vector machine (SVM) classifiers to distinguish known epilepsy-associated genes (Supp. File 1) from the rest of the genome based on their connectivity in a functional network. Each gene was assigned a functional association score, defined as the negative log of the average false positive rate, where the false positive rate reflects the proportion of unlabeled genes scoring above a given threshold. Full mathematical details are provided in Supplementary Methods. Training genes are provided in Supp. File 1.

### 2.2 Enrichment of High Functional Scoring Genes

A threshold of a functional score of 2 or higher was used as the threshold for a gene with high degree of relatedness to the epilepsy training genes. These genes were then used in a gene-set enrichment analysis. Significant gene sets were retrieved with the g:OST tool(14) to identify the enriched Gene Ontology (GO) biological process terms. We visualized these enrichments using tree maps with reviGO(15) to reduce the full set of enriched terms to a smaller number of semantically meaningful categories. A matrix reduction factor of 0.90 was used.

### 2.3 Loss of Function Tolerance and Functional Score

We retrieved the loss-of-function observed/expected upper bound fraction (LOEUF) score for each gene from the gnomAD v4.1 constraint metrics.(16) This score was calculated by taking the number of loss-of-function mutations observed divided by the upper boundary number of the predicted loss-of-function variants. We considered any gene with a LOEUF score less than 0.6 to be significantly intolerant to change as suggested by the gnomAD v4.1.(17,18) Bar graphs were generated using binned FS and LOEUF score.

### 2.4 Tractability and Functional Score

Tractability information was downloaded from Open Targets v24.09 tractability input table(19), which includes target status of each gene as “Approved Drug”, “Advanced Clinical”, “Phase 1 Clinical” from ChEMBL. We divided genes with a high functional score (FS > 2) into two groups based on the tractability information related to small molecule treatments, as these are the most likely to pass through the blood-brain barrier. One group contained genes that had an approved drug, or a compound in clinical trials while the other group contained genes that had no approved drug or small molecule treatment in clinical trial. These sets of genes were then used for a comparative enrichment analysis using the g:OST tool and enrichments of biological pathways between these two gene sets were compared and these pathways were reduced with a matrix reduction factor of 0.9 visualized with reviGO.(14,15)

### 2.5 Meta-analysis

To find genetic modifiers to epilepsy within mice we retrieved data from previous experiments on GeneNetwork using the search terms “epilepsy” and “seizure”.(12) These data included 127 quantitative measures of responses to multiple seizure-inducing stimuli, assayed in three major populations including BXD strains, AXBXA strains, and the hybrid mouse diversity panel (MDP). These measures were collated and uploaded to the Mouse Phenome Database (MPD)(20), and a GWAS meta-analysis was conducted using the METASOFT algorithm.(21) For a detailed breakdown of the measures and strains, see Supp. File 2.

### 2.6 Locus Identification

Meta-analysis output was downloaded from MPD and the p-value associated with each SNP was parsed from this output. We adopted a threshold of -log_10_(p)=15 to prioritize highly significant SNPs within this meta-analysis GWAS. Loci were defined as ±1Mb around a significant SNP, using the R function “findpeaks” from the “pracma” package to set the boundaries of each loci centered at a local peak. To resolve overlapping windows, loci with intersecting ranges were merged yielding 118 distinct loci, of varying sizes and number of genes.

### 2.7 Identification of SNP-Variant modifiers

Genes with a functional score of 2 or higher (FPR < 0.01) and within one of the 118 loci identified by meta-analysis were considered for further analysis. All SNPs within a gene of interest and 1kb flanking distance were retrieved from Genome MUSter.(22) To predict the deleterious effects of each SNP, we used the 91-mammals GERP conservation scores from the Ensembl “compara”, which give a score that indicates how conserved a position is across an alignment of the 91 mammals.(23) A high score indicates the site is highly conserved while a lower score indicates more variation is observed at that location. We filtered for SNPs with a GERP score of 2 or higher, which is the threshold for detecting SNPs with predicted deleterious effects suggested by the UCSC genome browser.(24) We then calculated the minimum hamming distance for each high-GERP SNP in a gene of interest and the SNPs that had a significant p-value in the meta-analysis to verify that the high-GERP SNPs had the same linkage pattern as the GWAS-identified SNP. SNPs with a hamming distance of 5 or less were considered to be linked to the statistically significant meta-analysis SNPs.

### 2.8 Identification of Modifier Genes

To identify SNPs that likely had a high impact within the evolutionary conserved positions we queried Ensembl with the remaining RSIDs and retrieved their functional annotations. These annotations were then manually reviewed to select SNPs with a larger predicted impact (i.e., splice, missense, and transcription factor binding variants). We focused on SNPs with significant alterations within regulatory regions. Additional literature reviews were conducted to narrow down the final candidate genes.

## 3 RESULTS

### 3.1 Functional network analysis of known human and mouse epilepsy genes reveals a shared interactome enriched for neural development, synaptic plasticity, and behavior

Hundreds of genes have been associated with rare and common forms of epilepsy using genetic mapping and candidate gene studies.(6) Emerging evidence suggests that these genes act within a complex system and converge on critical pathways that are common to many forms of epilepsy. (10) To identify the core shared pathways across diverse forms of epilepsy, we performed a network analysis to rank all genes in the genome for their functional association to a diverse set of known epilepsy genes. Using our previously published pipeline (25,26), we trained a machine learning classifier to classify genes as being epilepsy-related or not based on their connections to known epilepsy-related genes in human and mouse brain-specific gene interaction networks. As true positives, we pooled gene sets from the NHGRI GWAS Catalog, literature review, and Mouse Mine(6,27,28), resulting in a training set of 576 human genes and their respective mouse orthologs (Supp. File 1). The output of this classifier was a prioritized list of all genes in the genome, ranked by functional score for association with known epilepsy (see Supplemental Methods). We defined the epilepsy interactomes as being all genes with functional score > 2 (*i*.*e*., false positive rate < 1%)

To characterize the biological pathways enriched in the epilepsy interactomes, we used g:OST to identify the enriched Gene Ontology (GO) biological process terms.(14) We visualized these enrichments using tree maps to reduce the full set of enriched terms to a smaller number of semantically meaningful categories.(15) We found broadly concordant enrichments between species; both networks were enriched for processes including ion transport, nervous system development, synaptic signaling, and synapse organization (Fig. C), suggesting that the gene networks underlying epilepsy in humans and seizure phenotypes in mice are strongly conserved.

### 3.2 Functional score correlates with loss-of-function intolerance, suggesting that important epilepsy network genes are critical developmental hubs

Given the added value of genetic evidence for drug targets,(29) we sought to determine whether epilepsy interactomes were enriched for genes that are intolerant of loss-of-function mutations, as these are more likely to be disease-relevant genes.(30) To quantify this, we compared the functional scores in humans and mice to loss-of-function observed/expected upper bound fraction (LOEUF) scores from the gnomAD database, which uses human population genetic data to identify evolutionarily constrained sites.(16) As expected, genes with higher FS had significantly lower predicted LOEUF tolerance (Fig. 2 D), indicating that genes in the epilepsy interactome are under evolutionary constraint and that mutations in these genes are more likely to lead to disease.

**Fig. 2.**
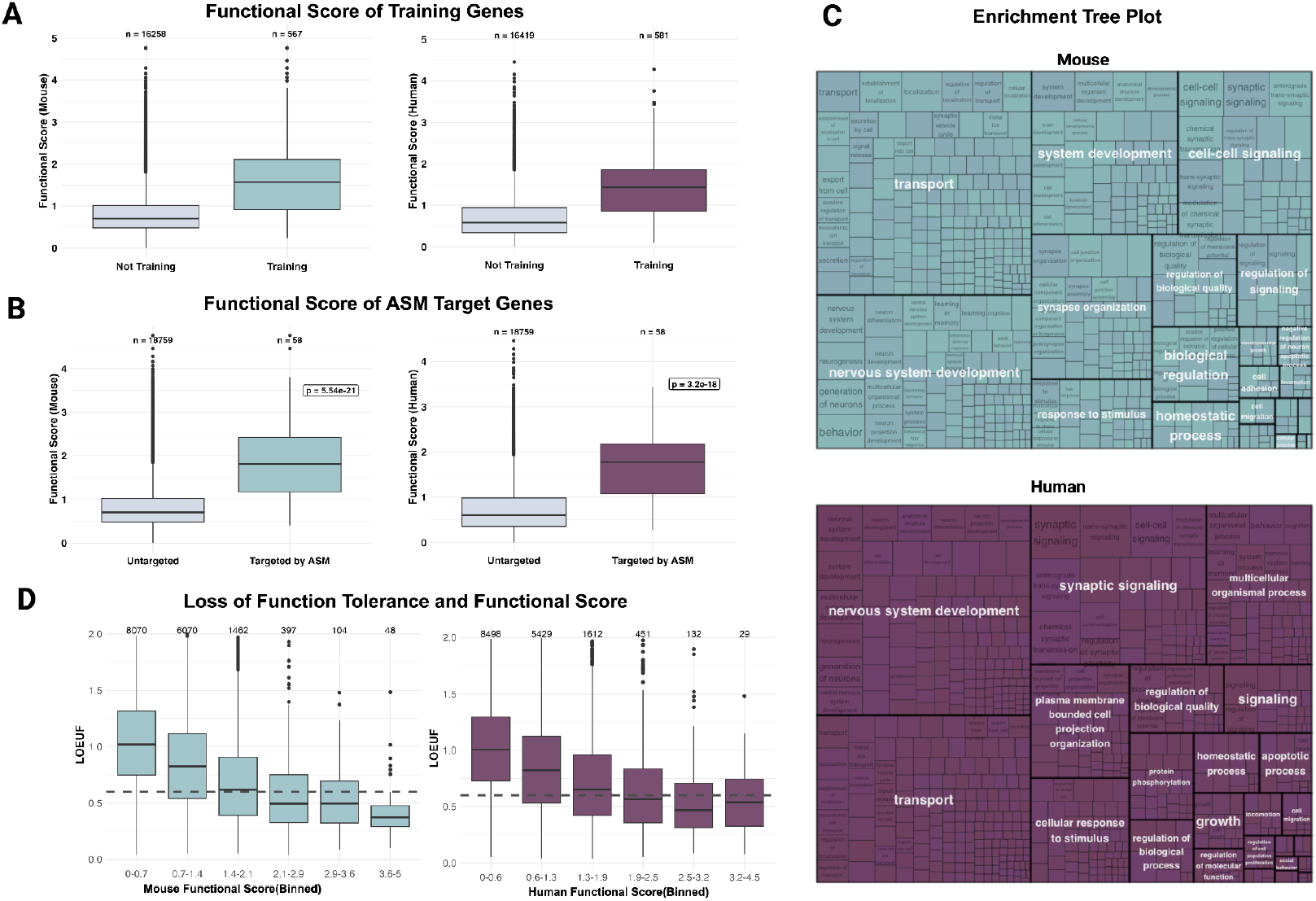
Functional network analysis of epilepsy genes in humans and mice. **(A)** As expected, genes used to train the SVM-ensemble had significantly higher functional scores than genes from outside the training set. Nevertheless, a subset of genes from outside the training set had high functional scores. **(B)** The functional scores of gene targets of existing ASMs was significantly higher in both humans and mice. **(C)** ReviGO tree maps of Gene Ontology enrichments of the top-ranked 300 genes from human and mouse interactomes revealed shared enrichments for neurodevelopment, synaptic plasticity, and membrane excitability processes. **(D)** Functional score was significantly negatively correlated to predicted loss-of-function (LOEUF) score in both the human and mouse interactomes.

### 3.3 Approved and late-stage-development drugs targeting high-ranking genes are enriched for membrane excitability and plasticity but not development

In order to characterize the currently exploited drug target landscape within the epilepsy interactome, we used the Open Targets Platform(19) to identify genes in the human epilepsy interactome (FS > 2) that are targeted by either an approved small molecule drug or a small molecule drug in Phase 3 clinical trials (Fig. 3; Supp. File 3). Overall, 78 out of 539 genes had an approved drug or a drug in development for some condition, including benzodiazepines, racetams, and topiramate. Next, we computed GO term enrichments for the set of targeted genes and the non-targeted genes. A scatter plot of the GO term p-values between these two sets revealed a difference in enrichment (Fig. 3A). Both the untargeted genes and the targeted genes were enriched for synaptic signalling, transport, and learning or memory (Fig. 3B). The targeted genes are highly enriched for processes related to ion transport, GABAergic synaptic transmission, and nerve impulse transmission (Fig. 3C). The non-targeted genes, on the other hand, are highly enriched for neurodevelopment and system development processes (Fig. 3D). This finding demonstrates that current ASM targets are well represented in the parts of the epilepsy interactome involved in membrane excitability, but poorly represented in the parts involved in neurodevelopment. Thus, there is a large collection of epilepsy-relevant biological processes that are outside of the current drug-development pipeline, but could potentially be exploited by future therapies.

**Fig. 3.**
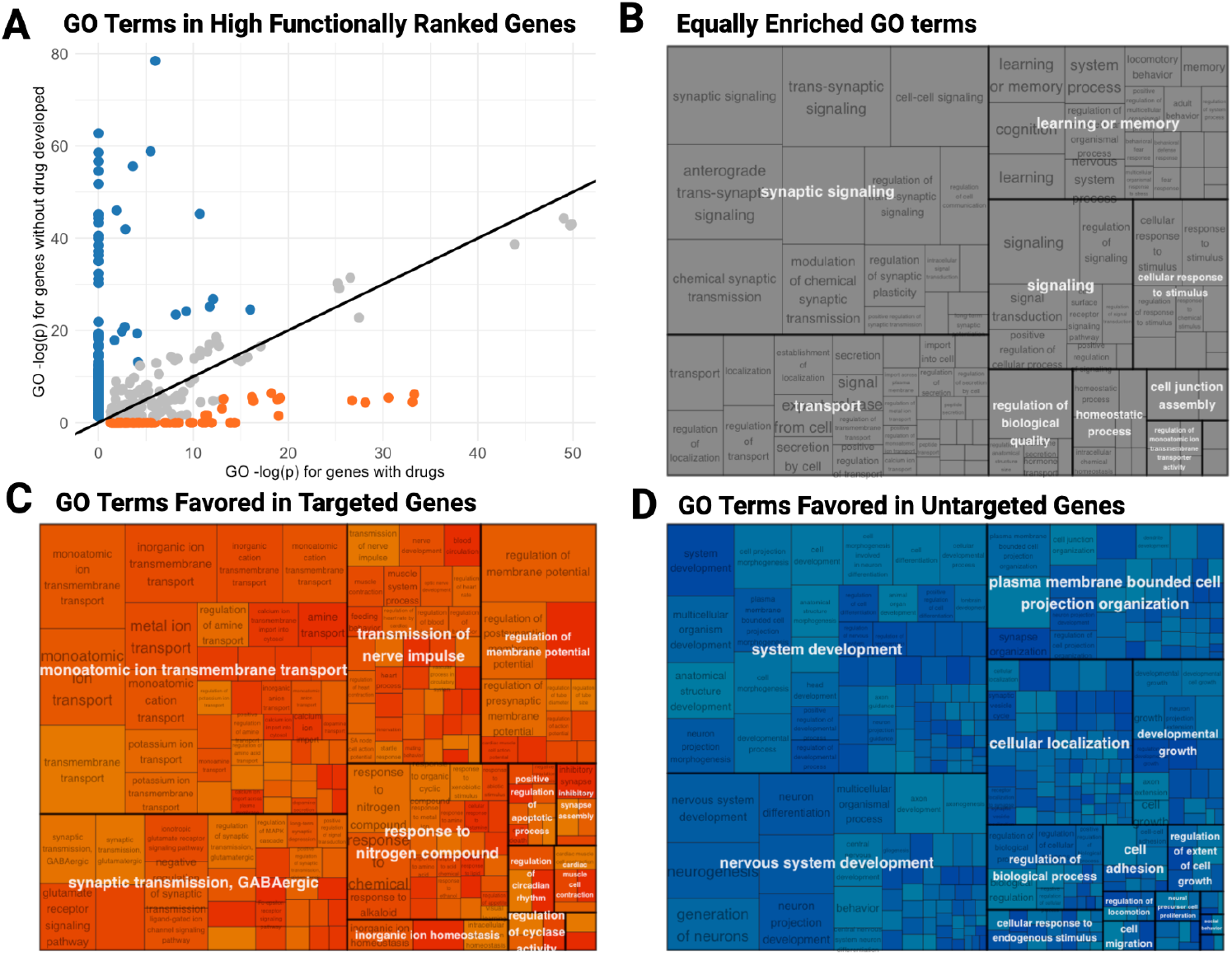
Comparison of Gene Ontology enrichments between ASM targets and untargeted genes within the human epilepsy interactome. **(A)** A scatter scatterplot of -log_10_(*p)* for enriched GO pathways in between the set of ASM targets within the human epilepsy interactome versus the untargeted genes in the human epilepsy interactome shows a set of shared enrichments, but also a set of divergent processes. **(B)** The ReviGO tree map of GO terms that are equally enriched in targeted and non-targeted genes show overlap in synoptic and plasticity processes underlying learning and memory. **(C)** The ReviGO tree map of GO terms that are preferentially enriched in ASM-target genes shows specific enrichments for genes involved in membrane excitability. **(C)** The ReviGO tree map of GO terms that are preferentially enriched in untargeted genes shows specific enrichments for processes involved in neurodevelopment.

### 3.4 Meta-analysis of 128 mouse seizure-related phenotypes reveals 118 novel modifier loci that are enriched for interactome genes involved in neurodevelopment and synaptic function

Given the widespread neurodevelopmental variability among inbred mouse strains,(12) which depends on plasticity mechanisms that are involved in early development to establish overall network architecture and adult cognition,(31,32) mice are an ideal model system to identify genetic modifiers of the epilepsy interactome that are outside the current target landscape. Furthermore, mouse experiments afford the opportunity to induce seizures in otherwise healthy backgrounds to identify resilience alleles that are independent of risk. To identify such alleles across multiple seizure-induction modalities, we performed a GWAS meta-analysis of 128 publicly available seizure-related phenotypes from the Gene Network database (12) that included the BXD and AXBXA recombinant inbred panels and the Mouse Diversity Panel (Table 1). Using the METASOFT analysis pipeline(21) implemented on the Mouse Phenome Database(20), we identified 118 distinct loci, represented on all 20 chromosomes (Fig. 4A). These findings demonstrate pervasive genetic influence on seizure-associated phenotypes throughout the mouse genome.

**Table 1.** Phenotypes and populations included in the GWAS meta-analysis

| Population | Measure | Induction Method |
| --- | --- | --- |
| BXD | Generalized Seizure Threshold | Fluorthyl |
|  | Kindling | Fluorthyl |
|  | Myoclonic Jerk Threshold | Fluorthyl |
|  | Incidence Severity Score | Audiogenic |
|  | Handling induced Convulsions | Ethanol Withdrawal |
|  | Handling induced Convulsions | Nitrous Oxide |
|  | Seizure Susceptibility | Atmospheric Pressure |
|  | Dosage Level | Cocaine, Pentylenetetrazol |
| MDP | Generalized Seizure Threshold | Fluorthyl |
|  | Kindling | Fluorthyl |
|  | Myoclonic Jerk Threshold | Fluorthyl |
|  | Maximal Electroshock Seizure Threshold | Electric Shock |
|  | CC50 for Psychomotor Seizure | Electrodes |
|  | CC50 for Maximal Tonic Hindlimb Extension Seizure | Electrode |
|  | ED50 against Maximal Tonic Hindlimb Extension Seizure | Electrode |
| AXBXA | Latency of Seizure | β-CCE |
|  | Anxiety related Behavior | Diazepam, Vehicle |

**Fig. 4.**
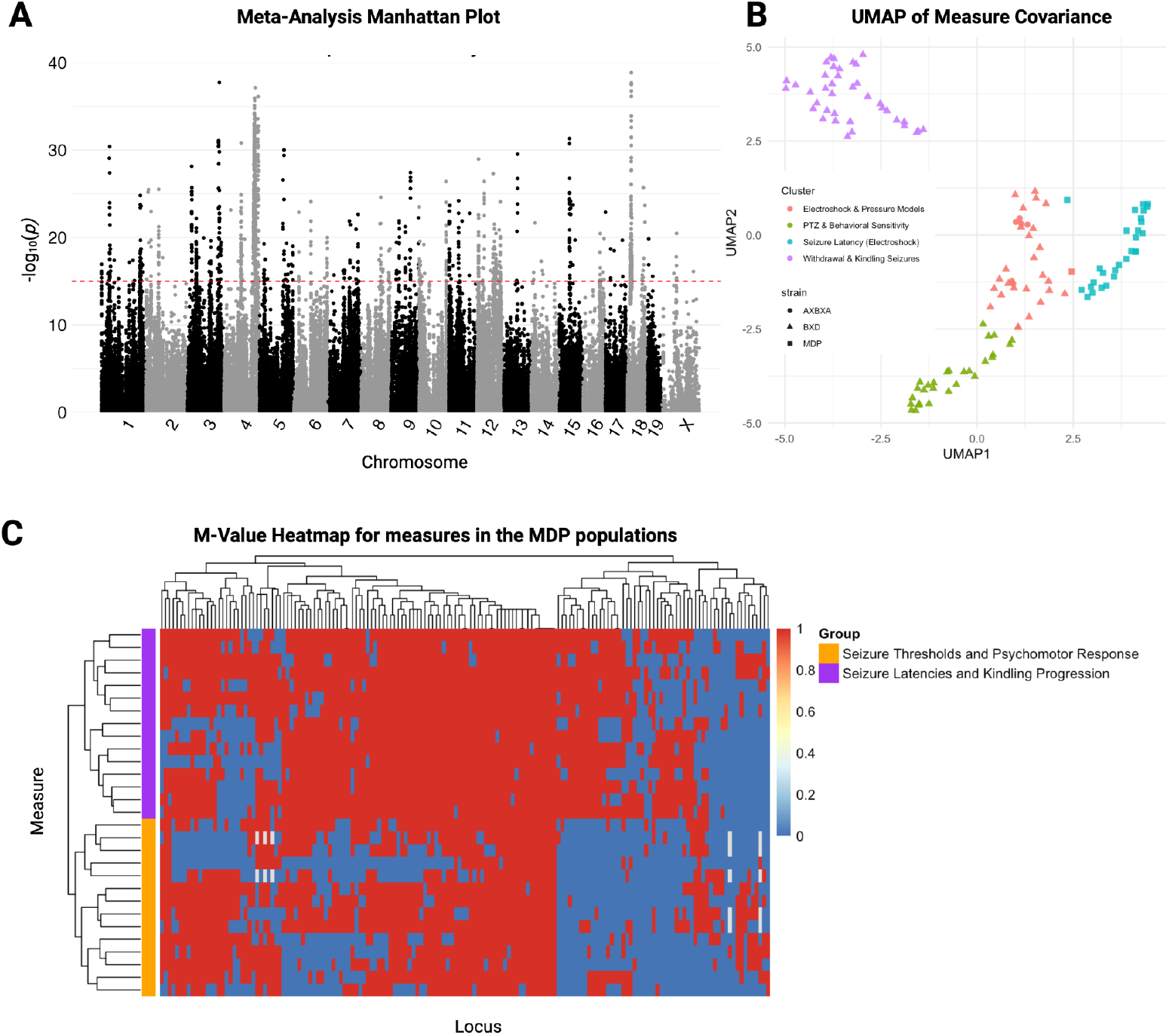
Mouse meta-analysis of 128 seizure-related phenotypes in mice. **(A)** The manhattan plot of meta-analysis results shows 118 distinct loci on all 20 mouse chromosomes (Y was excluded from analysis). **(B)** A UMAP dimension reduction analysis of the m-values from the meta-analysis shows clustering of genetic effects by study population and class of seizure phenotypes. **(C)** A heat map of the m-values for the phenotype measures studied in the Mouse Diversity Panel reveals two large groups of effect patterns, one related specifically to seizure latencies and kindling progression traits (purple traits, top) and another that influences both kindling progression and baseline seizure thresholds (purple and orange traits, bottom).

Given the large number of significant loci, we next sought to identify patterns in which loci were affecting which specific traits. Using an *m*-value cutoff of 0.9 (recommended by Han and Eskin(21)), we clustered traits and loci according to which loci had a significant influence on each trait (Fig. 4B). We found 4 trait clusters, with structure driven by the population in which they were studied (BXD, AXB, MDP) and according to the phenotype categories (method of induction and type of response). Nevertheless, we observe significant *m*-values from multiple populations and phenotypes from multiple populations for multiple SNPs, demonstrating the added value of pooling across populations. Within the cluster associated with the MDP studies, we identified two subclusters of loci, one cluster of loci that broadly influenced all traits and a second that was more specific to traits related to epileptogenesis after repeated exposures to flurothyl (Fig. 4D). Together, these results show that the putative modifier effects of the significant loci have complex, multidimensional effects on seizure phenotypes.

### 3.5 Integration of functional rankings, linkage analysis, and evolutionary conservation prioritize candidate variants within putative modifier genes

Variations in gene density and locus size led to large differences in the number of genes per locus. In total, these 118 loci contained 2822 genes, the majority of which are presumably only linked to a causal variant, rather than being seizure-modifier genes. Given the large number of genes in our 118 significant loci, we filtered the genes within QTLs down to potential seizure-modifier genes based on functional score (FS > 2), *i*.*e*., whether or not they are in the epilepsy interactome. Among the 2822 genes within the 118 QTLs, 168 genes met this criterion. The filtered list of 168 genes represents the positional candidate genes that have functional evidence suggesting they could be epilepsy modifiers (Supp. File 4).

To further filter the candidate gene list, we performed a comparative genomic analysis to identify potentially causal SNPs that could drive an effect targeting any of these genes. Using the Genome MUSter database(22) we retrieved the approximately 1 million mouse SNPs within these 168 genes. To estimate potential deleterious effects of these SNPs, we filtered by Genomic Evolutionary Rate Profiling (GERP) score from the 91-mammal GERP dataset from Ensembl.(23) This GERP score uses comparative genetics to estimate selection pressure on each position in the genome using multiple sequence alignments across 91 different mammals to define how often a position within the genome changes. Positions with lower variability across evolutionarily divergent species are predicted as more likely to be deleterious, which has been validated in multiple studies.(33–35) Using a GERP threshold of 2 (recommended by the UCSC genome browser),(24) we filtered down to 844 SNPs within the genes of interest (Supp. File 5). To determine which of the prioritized SNPs were linked to the markers identified by the meta-analysis, we filtered the 844 SNPs to those that segregated with the index SNPs using the minimum hamming distance between each candidate SNP and the index SNP in the corresponding locus. In total, 75 SNPs were found to have both a GERP score greater than 2 and a hamming distance less than or equal to 5 (Supp. file 6). These SNPs are high-quality candidate variants with convergent, multispecies evidence for modifier effects in epilepsy.

### 3.6 Bioinformatic prioritization predicts candidate variants that potentially drive altered gene expression

The above filtration steps considered gene function and evolutionary constraint to prioritize variants, but are unbiased with respect to the potential mechanisms of each SNPs effect (*e*.*g*., coding vs. non-coding, etc.). To identify potential mechanisms, we cross-referenced our filtered SNP list to the Ensembl database to retrieve functional annotations. Five genes –– *Ephb2, En2 Cadps2, Igsf21*, and *Cep170 ––* had SNPs within regulatory regions of their respective genes, potentially modifying local gene expression. Several of these genes have been linked to neurological abnormalities (Table 2), suggesting potentially novel roles as modifiers of epilepsy severity. No protein coding mutations were identified.

**Table 2.** Gene and SNP information for candidate modifiers of epilepsy. PV, parvalbumin; PC, pyramidal cell; HOX, homeobox; TLE, temporal lobe epilepsy.

| Gene | rsid | Regulatory Region | GERP | Pharos | Relevant Functions | PMID |
| --- | --- | --- | --- | --- | --- | --- |
| *Ephb2* | rs244989576 | Enhancer | 2.05 | Tchem | Synaptic structure, PV → PC connectivity, | 16626680, 39327008 |
| *En2* | rs46722611 | Enhancer | 4.05 | Tbio | HOX gene in brain development, GABAergic innervation | 19186208 |
| *Cadps2* | rs33763155 | Promotor | 2.73 | Tbio | Neurotrophin release, Number of PV cells | 17380209 |
| *Igsf21* | rs27612699 | Enhancer | 2.54 | Tdark | GABAergic synapse differentiation and growth | 38571813 |
| *Cep170* | rs32600451 | Promoter | 2.05 | Tbio | Centrosomal protein, In loci associated with TLE | 29904720 |

## 4 DISCUSSION

We performed a systems-level meta-analysis of epilepsy-associated genes in humans and mice to identify novel mechanisms that could be targeted by future epilepsy therapies. We identified a shared core gene network in both species and found that current drug-target genes are strongly enriched for membrane excitability but poorly enriched for neurodevelopmental processes, which makes sense given the critical importance of ion channels and neurotransmitter receptors in epilepsy pathophysiology, as well as the established preclinical discovery paradigm that emphasize acute seizures (36). The limitations of this paradigm, however, are widely known, especially in the context of developmental and epileptic encephalopathies (37). In this respect, genetic modifiers in diverse animal models provide essential insight into broader mechanisms of epilepsy pathophysiology (38). Our GWAS meta-analysis revealed 136 SNPs in 168 genes that had both a high conservation-based positional score (Fig. 4). Five of these SNPs––in the genes *Ephb2, Cadps2, Igsf21, En2*, and *Cep170* ––are annotated to regulatory regions, allowing us to bioinformatically prioritize these genes as high-confidence candidate modifiers of epilepsy. Importantly, these five genes have validated roles in neurodevelopmental and synaptic plasticity processes, including two with experimentally validated roles as epilepsy modifiers, that are outside the current drug target landscape for epilepsy.

*EPHB2/Ephb2* encodes ephrin type-B receptor 2 (EphB2), a member of the ephrin receptor family of receptor tyrosine kinases and is an established regulator of axonal growth and pathfinding. EphB2 participates in bidirectional cell signalling and has been studied in the context of cancer and autism spectrum disorder (ASD)(39). EphB2 is a negative regulator of PV to PC contact formation, suggesting that reduced activity could support increased inhibitory tone on excitatory cells. Mice with conditional *Ephb2* knockout in PV+ cells had an increased latency to onset of clonic seizures, decreased duration of seizure, and lower incidence of tonic–clonic seizures compared to wild type mice (40), supporting the hypothesis that reduced *Ephb2* expression may confer resilience to seizure-inducing stimuli. Three clinically approved tyrosine kinase inhibitors (bosutinib, dasatinib, and tivozanib) target EphB2 activity (41), but these are not highly selective to EphB2 making their effect on epilepsy difficult to predict (Supp. File.6). Nevertheless, as a member of a druggable receptor family with existing clinical therapeutics, and given its substantial role in pathways central to epilepsy, EphB2 emerges as a strong candidate target for epilepsy intervention. Given the availability of tyrosine kinase inhibitors that interact with EphB2, albeit non-selectively, there is a compelling rationale for further preclinical evaluation of these agents in epilepsy models.

Engrailed-2 (*EN2/En2*) is a homeobox-containing gene involved in the patterning of the central nervous system. In humans, *EN2* variants have been linked to neurodevelopmental abnormalities related to encephalic structural anomalies and altered *EN2* expression has been associated with ASD(42,43). *En2* knockout mice have autism-like behavioral phenotypes, along with increased susceptibility to kainic acid–induced seizures (44). Although *En2* is primarily classified as a developmental gene, its expression persists in the hippocampus and cerebral cortex of adult mice(44), suggesting potential for therapeutic intervention beyond early development. These findings suggest that En2 may serve as a dual-purpose target for both ASD and epilepsy, highlighting the need for therapeutic strategies that address overlapping neurodevelopmental pathways.

Calcium-dependent secretion activator 2 (*CADPS2*/*Cadps2)* encodes a protein involved in calcium-dependent vesicle secretion and regulates the release of neurotrophin-3 and brain-derived neurotrophic factor (BDNF). BDNF is crucial for the formation of synapses and neural circuits.(45) *Cadps2* knockout mice have significantly fewer PV-positive GABAergic interneurons in their cortex and hippocampus. Knockout mice also have a reduction in the spontaneous release of BDNF and display autistic-like behavior.(46) In humans *CADPS2* variants are causative for ASD and intellectual disability.(47) The reduction in PV-positive interneurons and BDNF release implicates CADPS2 in the maintenance of inhibitory tone, a critical factor in seizure susceptibility, suggesting its potential as a novel therapeutic target.

Immunoglobulin superfamily member 21 (*IGSF21*/*Igsf21*) is necessary for proper GABAergic synapse formation through interactions with neurexin-2α (Nrxn2α). Knockout of *Igsf21* leads to inhibition of GABAergic synapse transmission and sensorimotor deficits (48). Nrxn2α itself is strongly associated with multiple neurological disorders, including ASD and schizophrenia.(49) Given the specificity of *Igsf21* for GABAergic synapses, its modulation may offer a targeted approach to structurally restoring inhibition.

Centrosomal protein 170 (*CEP170/Cep170)* encodes for a centrosomal protein that is critical to cilia function and dynein-2 assembly. *CEP170* is located within a 2 Mb locus whose deletion has been associated with microcephaly, corpus callosum hypoplasia, and psychomotor retardation in humans. However, the presence of multiple plausible candidate genes within this region prevents definitive attribution of these phenotypes to *CEP170*.(50) Further functional studies are required to validate a potential role for *CEP170* in epilepsy.

## 5 CONCLUSION

Our results underscore the potential for novel therapeutic targets beyond traditional membrane excitability pathways, offering a broader landscape for anti-seizure medication (ASM) development. The identification of genes such as *EPHB2* and *EN2*, which have demonstrated influence on seizure susceptibility in knockout models, provides a compelling rationale for further preclinical and clinical investigation. Moreover, the involvement of these genes in neurodevelopmental and synaptic processes aligns with the complex etiology of treatment-resistant epilepsies, especially developmental and epileptic encephalopathies. While direct validation of these variants remains a future goal, our findings lay a foundational framework for translational research aimed at expanding the therapeutic arsenal for epilepsy. This work shows the power of multi-species, systems-level approaches to reveal potential therapeutic targets outside of membrane excitability that could advance ASM development.

## Acknowledgements

This research was funded by the National Science Foundation Grant #2244034.

